# Social isolation stress in adolescence, but not adulthood, produces hypersocial behavior in adult male and female C57BL/6J mice

**DOI:** 10.1101/2020.04.28.066704

**Authors:** Jean K. Rivera-Irizarry, Mary Jane Skelly, Kristen E. Pleil

## Abstract

Chronic stress during the developmental period of adolescence increases susceptibility to many neuropsychiatric diseases in adulthood, including anxiety, affective, and alcohol/substance use disorders. Preclinical rodent models of adolescent stress have produced varying results that are species, strain, sex, and laboratory-dependent. However, adolescent social isolation is a potent stressor in humans that has been reliably modeled in male rats, increasing adult anxiety-like and alcohol drinking behaviors, among others. In this study, we examined the generalizability and sex-dependence of this model in C57BL/6J mice, the most commonly used rodent strain in neuroscience research. We also performed a parallel study using social isolation in adulthood to understand the impact of adult social isolation on basal behavioral phenotypes. We found that six weeks of social isolation in adolescence beginning at postnatal day (PD) 28 produced a hypersocial phenotype in both male and female adults in multiple assays and a female-specific anxiolytic phenotype in the elevated plus maze, but it had no effects in other assays for avoidance behavior, fear conditioning, alcohol drinking, reward or aversion sensitivity, novel object exploration, or forced swim behavior in either sex. In contrast, social isolation in adulthood beginning at PD77 produced an anxiogenic phenotype in the light/dark box but had no effects on any other assays. Altogether, our results suggest that 1) adolescence is a critical period for social stress in C57BL/6J mice, producing aberrant social behavior in a sex-independent manner and 2) chronic individual housing in adulthood does not alter basal behavioral phenotypes that may confound interpretation of behavior following other laboratory manipulations.

## Introduction

Adolescence is a critical developmental period marked by increased reward seeking and impulsivity and the establishment of apposite social behaviors (Spear, 2004, Steinberg, 2004, Romer, 2010, Steinberg, 2010, Leshem, 2016). In humans, adolescence is associated with increased peer affiliation and separation from family (Noom et al., 1999, Keijsers et al., 2009, Eichelsheim et al., 2010). In rodents and other mammals, it is marked by heightened incidence of play behavior, altered social interactions, and increased exploration (Spear, 2004, Hawk et al., 2009, Trentacosta and Shaw, 2009, Walker et al., 2019). The quality and quantity of social interactions during adolescence have been linked to later-life behavioral outcomes in humans, including rates of drug and alcohol use and the formation of healthy social relationships (Bray et al., 2001, Kochenderfer-Ladd and Wardrop, 2001, Trentacosta and Shaw, 2009, Masten et al., 2012, Deutsch et al., 2015, Jager et al., 2015).

Adolescence is also marked by increased stress sensitivity, and chronic stress exposure during this period has been shown to alter brain structure and function (Paus, 2007, Eiland and Romeo, 2013). As peer interactions are especially important during adolescence (Steinberg, 2004, Jager et al., 2015), exposure to social stress may have particularly deleterious consequences on brain development and behavior (Casey et al., 2010, Platt et al., 2013, Burke et al., 2017). This increased stress sensitivity may partly explain why substance use disorders and many other psychiatric conditions frequently emerge during adolescence (Turner and Lloyd, 2004, Kessler et al., 2005, Kessler et al., 2007, Ernst and Fudge, 2009, Casey and Jones, 2010, Blakemore and Robbins, 2012). Understanding how adolescent social stress alters neurophysiology and behavior may prove crucial to treating stress-related disorders in adolescence and throughout later life.

Adolescent social isolation in rats has emerged as preclinical model that recapitulates many of the deleterious behavioral outcomes linked to chronic adolescent stress in humans (Lukkes et al., 2009b, Butler et al., 2016, Walker et al., 2019). In male rats, this paradigm has been shown to increase anxiety-like behavior and drug and ethanol intake and decrease fear memory extinction (McCool and Chappell, 2009, Whitaker et al., 2013, Butler et al., 2014a, Karkhanis et al., 2015, Skelly et al., 2015, Butler et al., 2016, Yorgason et al., 2016, Karkhanis et al., 2019), although these effects were not recapitulated in female rats (Butler et al., 2014b). Isolation during adolescence has also been linked to decreased social interaction in rats (Ferdman et al., 2007). Less is known about the effects of protracted adolescent isolation on these behaviors in mice, even though they are commonly used on neuroscience research, including studies that model human psychiatric conditions such as drug self-administration that requires individual housing (Becker and Ron, 2014). While some evidence suggests that isolation in adulthood is not stressful for mice (Hunt and Hambly, 2006), other work presents evidence to the contrary (Arakawa, 2018, Mumtaz et al., 2018, Manouze et al., 2019). The effects of isolation in adolescence are even less clear. Like humans, adolescent mice demonstrate a potentiated response to stress (Romeo et al., 2006). Although there are some reports that chronic social stress during adolescence increases depressive- and anxiety-like behaviors and drug self-administration in mice (Conrad and Winder, 2011, Lopez et al., 2011, Amiri et al., 2015), these results are variable and may be strain and sex-dependent (Arakawa, 2018, Mumtaz et al., 2018, Walker et al., 2019). C57BL/6J mice are commonly used in studies of alcohol self-administration (Rhodes et al., 2005, Melendez et al., 2006, Lyons et al., 2008, Yoneyama et al., 2008, Hwa et al., 2011, Mulligan et al., 2011) and as such are regularly singly housed for long periods of time. However, the lasting behavioral effects of social isolation (either in adolescence or adulthood) on escalated alcohol self-administration and anxiety-like behaviors in this strain have been variable (Lopez et al., 2011, Lopez and Laber, 2015, Huang et al., 2017, Caruso et al., 2018).

Here we evaluated the behavioral consequences of prolonged social isolation on behavior in male and female C57BL/6J mice and determined whether adolescence was a specific period of stress sensitivity. Following six weeks of social isolation in adolescence or adulthood, we measured anxiety, anhedonia, alcohol intake, reward and aversion sensitivity, fear memory formation and social behavior in adulthood. We found that social isolation produced few behavioral deficits overall, however this manipulation in adolescence led to aberrant social behavior in adulthood, marked by hyper-sociability and reduced avoidance behavior. Overall, these results suggest that single housing in adulthood does not robustly impact the basal behavioral state of C57BL/6J mice and that adolescence is a sensitive period for the effects of chronic social stress in this strain.

## Methods

### Animals

Male and female C57BL/6J mice were purchased from Jackson Laboratories (Bar Harbor, ME) at postnatal day (PD) 21 (for adolescent isolation experiment) or 63 (for adult isolation experiment) and housed on a 12 hr:12 hour light:dark cycle with lights off at 7:30 am and *ad libitum* access to food and water. One week after arrival, mice were randomly assigned to socially isolated (SI, one mouse per cage) or maintained in group housed (GH, five mice per cage) conditions for six weeks prior to behavioral testing. In the adolescent SI cohort, mice that were GH through adolescence were singly housed at PD77 for the duration of the study. In the adult SI cohort, GH mice remained in group-housed conditions. All experimental protocols were approved by the Institutional Animal Care and Use Committee at Weill Cornell Medicine in accordance with the guidelines of the NIH Guide for the Care and Use of Laboratory Animals.

### Behavioral Assays

Assays were conducted under 250 lux lighting conditions as previously described (Pleil et al., 2015, Crowley et al., 2016, Marcinkiewcz et al., 2016) and Panlab SMART 3.0 video tracking software was used to track and analyze behavior, unless otherwise described. Each behavioral apparatus was thoroughly cleaned with 70% ethanol prior to each trial. Timeline graphs illustrating the sequence of experiments conducted in the adolescent and adult isolation cohorts can be found in **Figures 1A** and **2A**, respectively.

**Figure 1.**
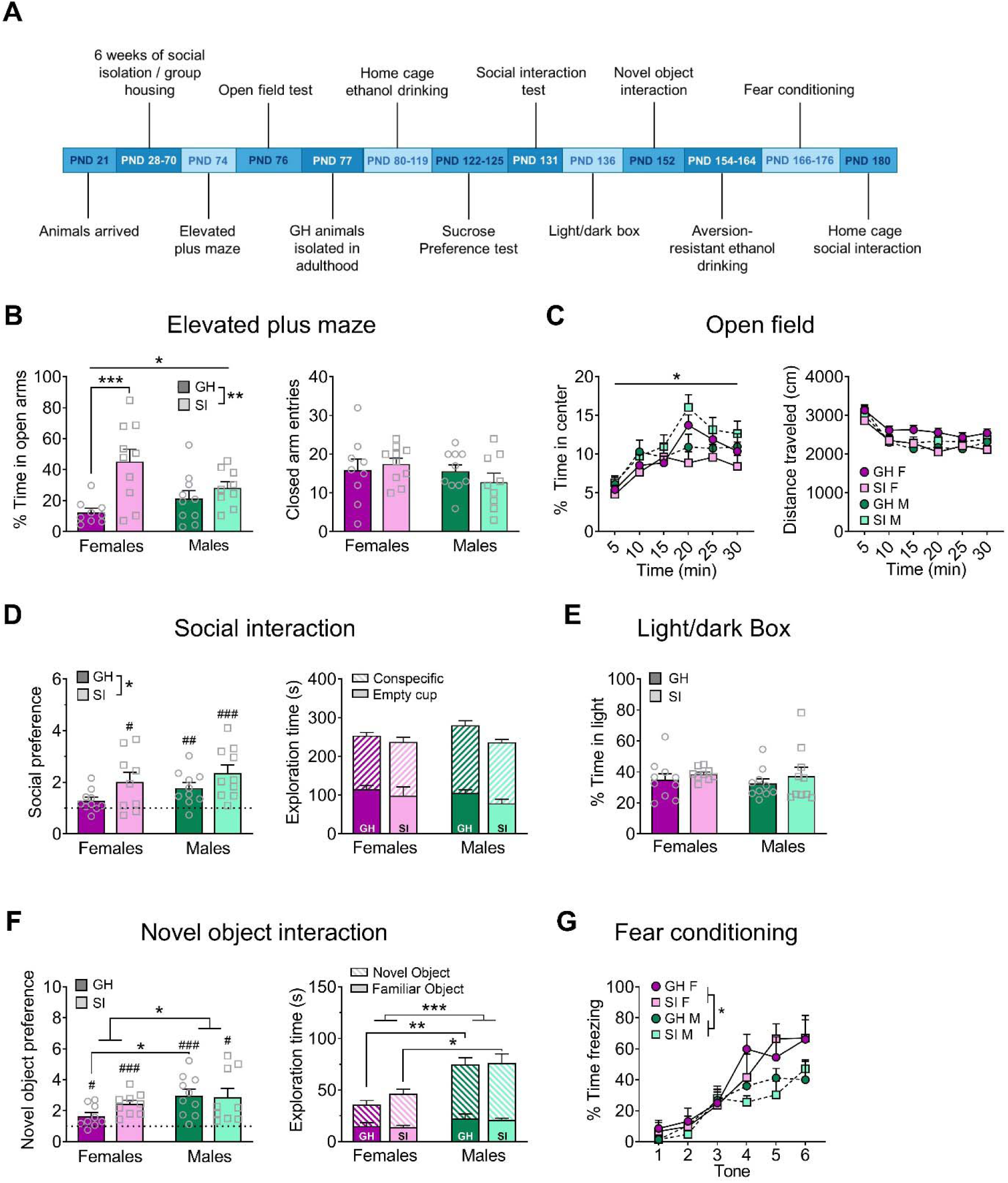
Adolescent social isolation behavior battery. (**A**) Experimental timeline. (**B**) In the elevated plus maze (EPM), adolescent social isolation (SI) increases the percent time spent exploring the open arms, an effect driven by females (left), without altering locomotor activity as measured by closed arm entries (right). (**C**) Adolescent SI oppositely affects the percent time spent exploring the center of an open field in males and females (left) but does not affect the distance traveled in this assay (right). (**D**) On the social interaction test, all but GH females display a significant preference for a novel social partner over an empty cup, and adolescent SI increases preference (left) without impacting total time spent exploring both objects (right). (**E**) Adolescent SI has no effect on anxiety-like behavior in the light/dark box. (**F**) All groups display a preference for a novel object over a familiar one, and this preference was greater in males than females but unaffected by adolescent SI (left). Total time spent exploring both objects is likewise increased in males compared to females (right). (**G**) Females display enhanced fear conditioning compared to males, but adolescent SI does not alter this measure. Data are expressed as means + SEM; \**p* < 0.05, \*\**p* < 0.01, \*\*\**p* < 0.001 between groups; ^#^*p* < 0.05, ^##^*p* < 0.01, ^###^*p* < 0.001 compared to null hypothesis of preference score = 1.

#### Elevated Plus Maze

The elevated plus maze (EPM) test was conducted in a plexiglass maze with two open and two closed arms (35 cm l × 5.5 cm w, with 15 cm h walls for closed arms) extending from a central platform (5.5 cm ×5.5 cm) elevated 50 cm above the floor. At the beginning of each trial, the mouse was placed in the center of the maze facing an open arm and movement was tracked continuously for five minutes. The total time spent on the open and closed arms of the assay and total number of open and closed arm entries (defined as placement of all four paws into the arm) were quantified. Percent time spent in the open arms of the assay was calculated to measure anxiety-like behavior, and closed arm entries were used as a measure of locomotion.

#### Open Field Test

The open field test was conducted in a plexiglass arena (50×50×34.5 cm) with a gray floor. The mouse was placed in one corner of the arena and allowed to explore freely for 30 minutes. Total time spent in the center of the maze (defined as having all four paws in the 25 cm x 25 cm area in the center of the arena) and periphery were quantified to calculate percent center time. The total distance traveled in the maze (cm) was used to measure locomotion, and percent time in the center of the maze was used to assess anxiety-like behavior.

#### Light/Dark Box

The light/dark box assay was conducted in a rectangular box divided into two equal compartments (20 cm l × 40 cm w × 34.5 cm h), one dark with a closed lid and the other with an open top and illuminated by two 60 W bulbs placed 30 cm above the box. The two compartments were separated by a divider with a 6 cm x 6 cm cut out passageway at floor level. At the beginning of each trial, the mouse was placed in a corner of the light compartment and allowed to move freely between the two compartments for 10 minutes. The number of light box entries and total time spent in the light compartment as compared to the dark compartment were used to assess anxiety-like behavior.

#### Social Interaction Test

The social interaction test was conducted in three 10-minute phases in an open plexiglass arena (50 cm × 50 cm × 34.5 cm), and mice could explore freely during each phase. Between each testing phase, the experimental mouse was briefly placed in a holding cage while the experimenter altered the arena settings as follows: phase 1: empty arena; phase 2: two empty wire mesh cups (diameter 4”, height 4”) located at opposite corners of the arena 10 cm from each wall; phase 3: a novel age- and sex-matched mouse of the same strain was placed inside one of the two cups, while the other cup remained empty. The placement of the cups and social partner were pseudorandom and counterbalanced. Interaction zones for each cup were defined as encompassing a 5 cm radius around the center of the cup, and the ratio of interaction time with the social partner versus the empty cup during phase 3 was used to determine a social preference score.

#### Novel Object Interaction

The novel object interaction assay was conducted under the same conditions and using the same analyses as the social interaction test (see above) but using objects, in order to assess whether effects observed in novel social partner preference could be generalized to a non-social novel object. The objects used in this experiment included plastic cuboids with orange color (3 cm × 3 cm × 6 cm) and half-sphered plastic cylinders with a blue color of the same dimensions, as described in previous publications (Lueptow, 2017, Tian et al., 2019); these objects were determined to be of equal interest to C57BL/6J mice in pilot testing. The objects were affixed to the floor of the arena during behavioral testing, which proceeded as follows: phase 1: empty arena; phase 2: two versions of the same object located at opposite corners of the arena 10 cm from each wall; phase 3: a novel object replaced one of the two familiar objects in the arena. The ratio of interaction time with the novel versus familiar object during phase 3 was used as a novel object preference score.

#### Fear Conditioning

Fear conditioning was performed in an operant box with a stainless-steel grid floor within a sound-attenuating chamber (Colbourn Instruments, Allentown, PA). The mouse was placed in the chamber at the beginning of the test, and following a five min habituation period received six pairings of a 30 second, 80 dB tone (conditioned stimulus, CS) co-terminating with a 2 second, 0.5 mA foot shock (unconditioned stimulus, US) separated by pseudorandom intra-interval times (from 31-119 seconds, with an average ITI of 75.5 seconds). Video tracking and FreezeFrame software (Colbourn Instruments, Allentown, PA) were used to assess freezing behavior during the 28 second period preceding the shock across tone/shock presentations.

#### Home Cage Ethanol Drinking

We used a modified version of the standard Drinking in the Dark (DID) binge ethanol drinking paradigm (mDID) to assess binge ethanol intake under limited-access conditions as well as 24-hour preference for ethanol over water. Mice were singly housed for several days prior to the first ethanol presentation. For each mDID cycle, the home cage water bottle was replaced with a bottle containing 20% (cycles 1-4) or 30% (cycles 5-6) ethanol for two hours beginning three hours into the dark cycle for three days. On day 4, two bottles (one containing ethanol solution, one containing water) were placed in the cage for 24 hours (bottles were weighted after 2 hours, 4 hours, and 24 hours of access). Bottle weights were used to calculate ethanol and water consumption daily (normalized to bodyweight) and 24 hr ethanol preference on day 4, calculated as the ratio of the volume of liquid consumed from the ethanol bottle to the water bottle.

#### Aversion-Resistant Ethanol Drinking

Consumption and preference of quinine-adulterated ethanol over water in a two-bottle choice home cage assay was measured to evaluate aversion-resistant ethanol drinking behavior. Mice received 4 hours of access to two bottles, one containing 20% ethanol adulterated with 100 μM (Days 1 and 2) or 250 μM (Day 3) quinine hemisulfate (Sigma-Aldrich, St. Louis, MO) and the other containing water. Bottle placement was pseudorandom and switched daily, and consumption and preference were measured as described for mDID.

#### Sucrose Preference Test

A similar procedure to that described above was used to evaluate consumption and preference for 1% (w/v) sucrose solution versus water, except that mice were given access to the sucrose and water bottles for 24 hours per day. Intake and preference were measured every 24 hours for four consecutive days. For all drinking experiments, empty “dummy” cages on the same rack as housed behavior mice received the same ethanol, sucrose or water bottle replacement, and consumption was adjusted for leak from dummy bottles and normalized to bodyweight.

#### Home Cage Social Interaction

Home cage social interaction with a novel same-sex conspecific mouse was conducted in the experimental mouse’s home cage (28 cm × 18 cm × 12.5 cm). The novel mouse was placed into the cage and overhead video was used to record behavior for five minutes. An experimenter blind to condition hand-scored discrete behaviors performed by the experimental mouse, including the number and duration of total, head-to-head, and head-to-tail social interactions, as well as digging and climbing bouts.

### Statistical Analysis

Statistical analyses were conducted using GraphPad Prism 8 software. Distributions of data within group were analyzed for normality, and outliers were identified using Q-Q plots and confirmed by the Rout method (Q = 0.5%); when an individual mouse’s behavior was identified as an outlier for at least half of the reported dependent measures for an assay, it was excluded from analysis for that assay. Two-way analysis of variance (ANOVA) was used to assess the effects of housing condition and sex on behavior in the elevated plus maze, open field test (adult cohort), novel object test, light/dark box, and social interaction assays. Two-way repeated measures ANOVA (RM ANOVA) was used to assess the effects of housing condition on home-cage drinking behaviors within sex. Three-way RM ANOVA was used to assess the freezing across consecutive tone/shock pairings in the fear conditioning assay and behavior in the open field test across time (adolescent cohort). Equal variance across time was not assumed in RM-ANOVAs with three or more repeated measures, and a Greenhouse-Geisser correction of degrees of freedom was used. Significant effects in all ANOVAs were followed up with post-hoc two-tailed t-tests corrected for multiple comparisons using the Holm-Sidak method, and adjusted *p* values are presented. Alpha values of 0.05 were used throughout all analyses, and data are presented as mean + SEM.

## Results

### Elevated Plus Maze

Following six weeks of adolescent SI or GH conditions, mice underwent testing in the EPM to assess differences in anxiety-like behavior (**Figure 1B**; GH females n = 9, GH males n = 10, SI females n = 10, SI males n = 9). A two-way RM ANOVA comparing the percent time spent on the open arms revealed a main effect of housing condition (*F*_(1,34)_ = 12.78, *p* = 0.001) but no main effect of sex (*F*_(1,34)_ = 0.53, *p* = 0.472) and a significant interaction between sex and housing condition (*F*_(1,34)_ = 0.41, *p* = 0.026). Post-hoc analysis showed that this effect was driven by females, as SI females spent significantly more time on the open arms than their GH counterparts (*t*_(34)_ = 4.17, adjusted *p* = 0.0004), while SI males did not (adjusted *p* > 0.05). A two-way RM ANOVA on the number of closed arm entries revealed no effects of housing (*F*_(1,34)_ = 0.08, *p* = 0.776) or sex (*F*_(1,34)_ = 1.41, *p* = 0.776), nor a sex by housing condition interaction (*F*_(1,34)_ = 1.10, *p* = 0.301), suggesting that the increased open arm exploration in SI females was not due to a general increase in locomotion.

In contrast, social isolation during adulthood did not alter anxiety-like behavior on the EPM (**Figure 2B**). A two-way RM ANOVA revealed a main effect of sex (*F*_(1,34)_ = 6.66, *p* = 0.014) but no main effect of housing condition (*F*_(1,34)_ = 0.01, *p* = 0.928) nor a sex by housing condition interaction (*F*_(1,34)_ = 0.10, *p* = 0.753). Despite this significant main effect of sex in the omnibus test, post-hoc analysis did not reveal any significant differences between males and females (adjusted *p* > 0.05). A two-way RM ANOVA on the number of closed arm entries revealed no effects of sex (*F*_(1, 34)_ = 3.17, *p* = 0.084) or housing condition (*F*_(1, 34)_ = 0.33, *p* = 0.569), nor was there a significant interaction between these factors (*F*_(1, 34)_ = 1.95, *p* = 0.171).

**Figure 2.**
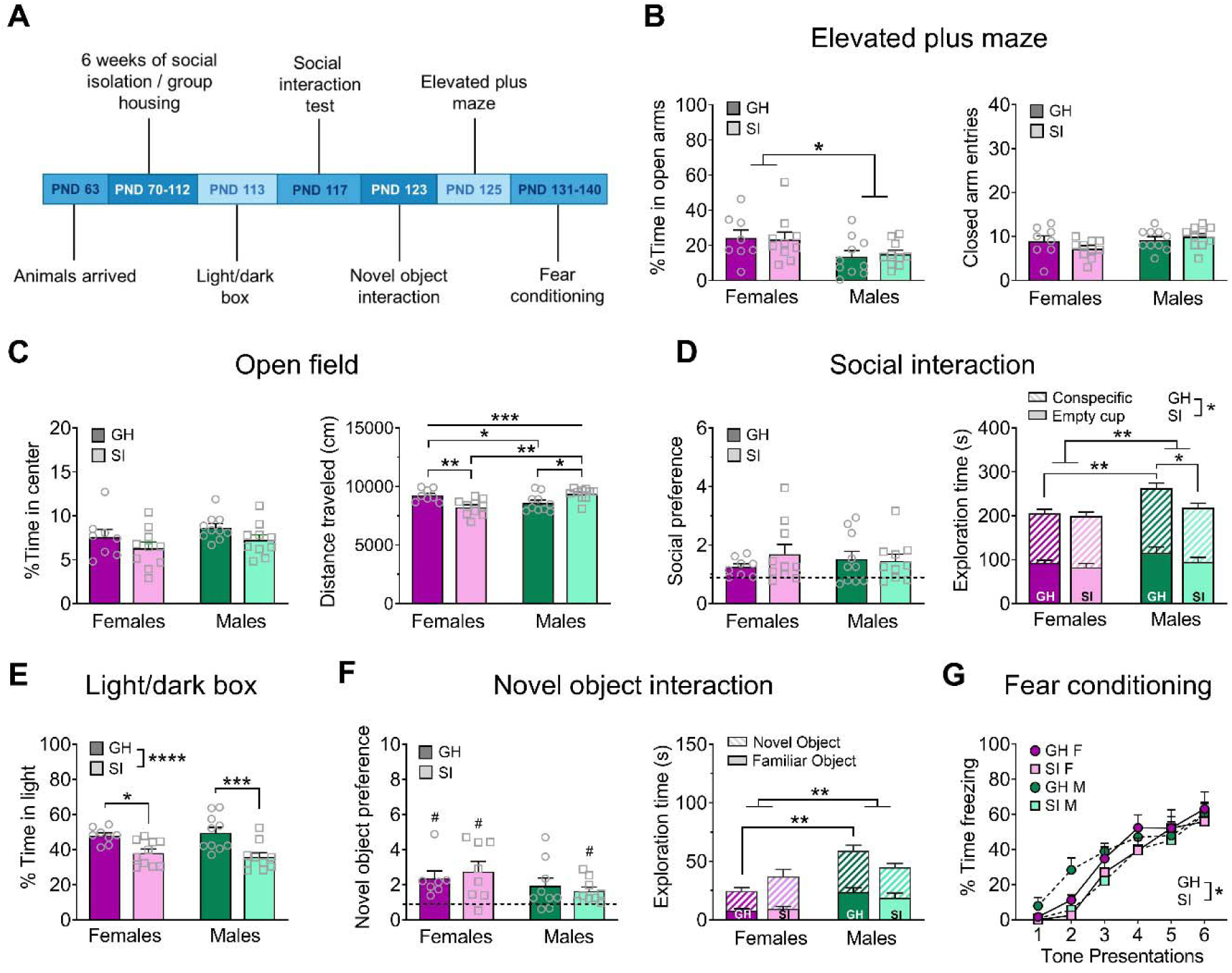
Adult isolation behavior battery. (**A**) Experimental timeline. (**B**) Females spend more time exploring the open arms of the EPM, but adult SI does not influence this measure (left); there are no difference in general locomotor behavior, measured by the number of entries into the closed arms (right). (**C**) There are no effects of sex or adult SI on the percent time spent exploring the center of the OF (left), however there is a sex-dependent effect of adult SI on the total distance traveled in the OF (right). (**D**) Adult SI does not alter preference for a novel social partner over an empty cup in the social interaction test (left) but does decrease total time spent interacting with the social partner and empty cup, an effect driven by males (right). GH males also spend more total time exploring both objects compared to GH females. (**E**) Adult SI decreases the percent time spent exploring the light side of the light/dark box in both males and females. (**F**) In the novel object interaction test, all groups except GH males display a preference for a novel vs. familiar object (left), however this is driven by greater overall interaction time with both objects in males (right). (**G**) Adult SI mice show delayed fear acquisition compared to GH mice. Data are expressed as means + SEM; \**p* < 0.05, \*\**p* < 0.01, \*\*\**p* < 0.001, \*\*\*\**p* < 0.001 between groups; ^#^*p* < 0.05 compared to null hypothesis of preference score = 1.

### Open Field Test

To further investigate the impact of adolescent social isolation on anxiety-like and locomotor behavior in early adulthood, we next compared open field exploration in GH and SI female and male mice (**Figure 1C**; n = 10 per group). A three-way RM ANOVA comparing the impact of sex and adolescent housing condition on the percent time spent in the center of an open field across time (30 minutes total, broken into 5 minute intervals) revealed a significant main effect of time (*F*_(5,180)_ = 18.63, *p* < 0.0001) but no effects of sex (*F*_(1,36)_ = 3.20, *p* = 0.082) or housing condition (*F*_(1,36)_ = 0.001, *p* = 0.971). No significant interactions were identified between time and sex (*F*_(5,180)_ = 0.31, *p* = 0.906), time and housing condition (*F*_(5,180)_ = 0.31, *p* = 0.904), or sex and housing condition (*F*_(1,36)_ = 3.35, *p* = 0.075). While there was a significant three-way time by sex by housing condition interaction (*F*_(5,180)_ = 2.94, *p* = 0.014), post-hoc analysis did not reveal any significant comparisons (adjusted *ps* > 0.05). A three-way RM ANOVA comparing the total distance traveled in the open field across these time points revealed a significant main effect of time (*F*_(5,180)_ = 57.65, *p* < 0.0001) but no main effects of sex (*F*_(1,36)_ = 0.53, *p* = 0.473) or housing condition (*F*_(1,36)_ = 1.66, *p* = 0.205). There was an interaction between time and sex (*F*_(5,180)_ = 2.41, *p* = 0.038) but no significant interaction between time and housing condition (*F*_(5,180)_ = 0.85, *p* = 0.516) or sex and housing condition (*F*_(1,36)_ = 4.01, *p* = 0.052), and no three-way interaction between time, sex, and housing condition (*F*_(5,180)_ = 1.57, *p* = 0.171). Post-hoc analysis did not reveal any significant differences between sexes at any time point, however (adjusted *p* > 0.05).

In the adult SI cohort, we used a 10 min open field test (**Figure 2C**; GH females n = 8, GH males n = 10, SI females n = 10, SI males n = 10). A two-way RM ANOVA comparing the percent time in the center of this assay did not reveal a main effect of sex (*F*_(1,34)_ = 2.29, *p* = 0.139) or housing condition (*F*_(1,34)_ = 4.07, *p* = 0.051), and the interaction between these variables also failed to achieve significance (*F*_(1,34)_ = 0.01, *p* = 0.931). Interestingly, a two-way RM ANOVA comparing the total distance traveled during this five minute assay did not reveal main effects of sex (*F*_(1,34)_ = 1.38, *p* = 0.248) or housing condition (*F*_(1,34)_ = 0.41, *p* = 0.526) but did reveal a significant interaction between these factors (*F*_(1,34)_ = 18.72, *p* = 0.001). Post-hoc comparisons revealed that GH females traveled a greater distance than their SI counterparts (*t*_(34)_ = 3.41, adjusted *p* = 0.005) while GH males traveled significantly less distance in this assay than SI males (*t*_(34)_ = 2.69, adjusted *p* = 0.021). Furthermore, the total distance traveled was higher in GH females than GH males (*t*_(34)_ = 2.17, adjusted *p* = 0.037), and higher in SI males than SI females (*t*_(34)_ = 4.01, adjusted *p* = 0.001).

### Social Interaction Test

To determine whether chronic social isolation during adolescence effects adult social behavior, mice in the adolescent SI cohort underwent a social interaction test (**Figure 1D**; GH females n = 10, GH males n = 10, SI females n = 9, SI males n = 10). Male and female mice reared in isolation, as well as GH males, demonstrated a significant preference for a social partner as compared to an empty cup (one-sample *t*-tests; GH males, *t*_(9)_ = 2.15, *p* = 0.004; SI females, *t*_(8)_ = 2.69, *p* = 0.027; SI males, *t*_(9)_ = 4.40, *p* = 0.001); however adolescent GH females did not demonstrate this social preference (*t*_(9)_ = 2.15, *p* = 0.060). Interestingly, a two-way RM ANOVA analyzing preference for a social partner over a non-social object revealed a significant main effect of housing condition (*F*_(1, 35)_ = 5.98, *p* = 0.019) but no main effect of sex (*F*_(1, 35)_ = 2.49, *p* = 0.123) or interaction between these factors (*F*_(1, 35)_ = 0.07, *p* = 0.787). However, post-hoc analysis did not reveal any significant differences in social preference between GH and SI animals of either sex (adjusted *p* > 0.05). A two-way RM ANOVA assessing general activity in this assay, as measured by combining the total time spent exploring both a social partner and a non-social empty cup, revealed no significant differences between groups (main effect of sex: *F*_(1, 35)_ = 0.50, *p* = 0.484; main effect of housing condition: *F*_(1, 35)_ = 2.69, *p* = 0.110; sex by housing condition interaction: *F*_(1, 35)_ = 0.55, *p* = 0.462).

In the adult SI cohort (**Figure 2D**; GH females n = 8, GH males n = 10, SI females n = 10, SI males n = 10), no group demonstrated a reliable preference for a social partner over an empty cup (one-sample *t*-tests; GH females: *t*_(7)_ = 2.23, *p* = 0.060; GH males: *t*_(9)_ = 1.87, *p* = 0.094; SI females: *t*_(9)_ = 2.10, *p* = 0.065; SI males: *t*_(9)_ = 2.05, *p* = 0.070). A two-way RM ANOVA did not reveal significant differences in social preference between groups (main effect of sex: *F*_(1, 29)_ = 3.15, *p* = 0.086; main effect of housing condition: *F*_(1, 29)_ = 0.02, *p* = 0.896; sex by housing condition interaction: *F*_(1, 29)_ = 0.59, *p* = 0.448). A two-way RM ANOVA comparing the total combined time spent exploring both the non-social object (empty cup) and social partner revealed significant main effects of sex (*F*_(1, 34)_ = 10.04, *p* = 0.003) and housing condition (*F*_(1, 34)_ = 4.32, *p* = 0.045), but there was no interaction between these factors (*F*_(1, 34)_ = 2.51, *p* = 0.122). Follow-up post-hoc analyses revealed that GH males spent more combined time exploring a social partner and empty cup than GH females (*t*_(34)_ = 3.27, adjusted *p* = 0.010) and SI males (*t*_(34)_ = 2.67, adjusted *p* = 0.034).

### Light/Dark Box

A two-way RM ANOVA did not reveal any effects of adolescent social isolation or sex (**Figure 1E**; n = 10 per group) on the percent time spent in the light side of a light/dark box (main effect of sex: *F*_(1, 35)_ = 0.21, *p* = 0.646; main effect of housing condition: *F*_(1, 35)_ = 1.21, *p* = 0.279; sex by housing condition interaction: *F*_(1, 35)_ = 0.023, *p* = 0.879). A two-way RM ANOVA comparing the effects of six weeks of adult social isolation versus group housing conditions on behavior in the light/dark box in males and females (**Figure 2E**; GH females n = 8, GH males n = 10, SI females n = 10, SI males n = 10) revealed a significant main effect of housing condition (*F*_(1, 34)_ = 21.78, *p* < 0.0001), but no main effect of sex (*F*_(1, 34)_ = 0.020, *p* = 0.886) or significant interaction between these variables (*F*_(1, 34)_ = 0.550, *p* = 0.463). Post-hoc analysis revealed that GH animals spent significantly more time in the light compartment of the light/dark box than their SI counterparts (GH males versus SI males: *t*_(34)_ = 3.94, adjusted *p* = 0.0008; GH females vs SI females: *t*_(34)_ = 2.70, adjusted *p* = 0.011).

### Novel Object Interaction

To determine whether the increased social exploration observed following adolescent social isolation could be generalized to non-social contexts, we performed a novel object interaction task designed similarly to the social interaction task described above (**Figure 1F**). GH females (n = 9) demonstrated a preference for a novel object over a familiar object (one-sample *t*-test, *t*_(8)_ = 2.71, *p* = 0.026), as did GH males (n = 10; *t*_(9)_ = 4.83, *p* = 0.0009), SI females (n = 9; *t*_(8)_ = 6.02, *p* = 0.0003), and SI males (n = 9; *t*_(8)_ = 3.29, *p* = 0.011). A two-way ANOVA comparing novel object preference across groups revealed a significant main effect of sex (*F*_(1, 33)_ = 5.20, *p* = 0.029) but no main effect of housing condition (*F*_(1, 33)_ = 0.766, *p* = 0.387) or significant interaction between these factors (*F*_(1, 33)_ = 1.31, *p* = 0.261). Post-hoc analysis revealed that GH males exhibited a significantly increased novel object preference as compared to GH females (*t*_(33)_ = 2.45, adjusted *p* = 0.039). To assess general exploratory behavior in this assay, we compared the total time that animals in each group spent exploring both the novel plus familiar objects in this assay. A two-way ANOVA revealed a significant main effect of sex (*F*_(1, 33)_ = 17.91, *p* = 0.0002), but no main effect of housing condition (*F*_(1, 33)_ = 0.54, *p* = 0.469) or interaction between these factors (*F*_(1, 33)_ = 0.36, *p* = 0.553). Post-hoc analysis revealed that GH females spent significantly less time exploring the novel and familiar objects than GH males (*t*_(33)_ = 3.46, adjusted *p* = 0.002). Consistent with this, SI females also spent less time exploring these objects that SI males (*t*_(33)_ = 2.54, adjusted *p* = 0.016). Altogether, these results suggest that while there are sex differences in the preference for and exploration of novel objects over familiar, adolescent social isolation had no effect on this task. In contrast, adolescent social isolation increased preference for a social partner, suggesting that its effects were specific to a social context.

In the adult SI cohort (**Figure 2F**), GH females displayed a significant preference for the novel object (n = 7; *t*_(6)_ = 3.13, *p* = 0.026), as did SI females (n = 8; *t*_(7)_ = 3.07, *p* = 0.017) and SI males (n = 9; *t*_(8)_ = 2.84, *p* = 0.022), but not GH males (n = 9; *t*_(8)_ = 1.99, *p* = 0.082). A two-way ANOVA comparing novel object preference across groups revealed no significant differences between groups (main effect of sex: *F*_(1, 29)_ = 3.15, *p* = 0.896; main effect of housing condition: *F*_(1, 29)_ = 0.017, *p* = 0.896; sex by housing condition interaction: *F*_(1, 29)_ = 0.59, *p* = 0.448). A two-way ANOVA comparing the total combined time spent exploring the novel and familiar objects revealed a significant main effect of sex (*F*_(1, 29)_ = 10.64, *p* = 0.002), but no main effect of housing condition (*F*_(1, 29)_ = 0.019, *p* = 0.890) or sex by housing interaction (*F*_(1, 29)_ = 4.13, *p* = 0.051). Post-hoc tests revealed that GH males spent significantly more combined time exploring a social partner and novel object than GH females (*t*_(29)_ = 3.68, adjusted *p* = 0.002)

### Fear Conditioning

We next assessed whether adolescent social isolation impacts fear learning by measuring acquisition of freezing behavior in response to a foot shock-paired tone (assessed by freezing during tone presentation across six consecutive tone/shock pairings; **Figure 1G**). A three-way RM ANOVA was used to measure the effects of sex and adolescent housing condition on freezing behavior across time (GH females n = 5, SI females n = 4, GH males n = 5, SI males n = 5). This test revealed a significant main effect of time, as expected (*F*_(3,045, 45)_ = 34.28, *p* < 0.0001). A main effect of sex also emerged (*F*_(1, 15)_ = 5.36, *p* = 0.035) as well as a significant time by sex interaction (*F*_(5, 75)_ = 2.68, *p* = 0.027). There was no significant main effect of housing condition (*F*_(1, 15)_ = 0.23, *p* = 0.638), time by housing condition interaction (*F*_(5, 75)_ = 0.80, *p* = 0.550), sex by housing condition interaction (*F*_(1, 15)_ = 0.010, *p* = 0.919), or time by sex by housing condition interaction (*F*_(5, 75)_ = 0.63, *p* = 0.680). Post-hoc comparisons did not reveal any significant sex-dependent differences at any time point, however (adjusted *p* > 0.05).

We also assessed fear conditioning in the adult SI cohort (**Figure 2G**; GH female n = 8, GH male n = 10, SI female n = 10, SI male n = 10). A three-way RM ANOVA revealed a main effect of time (*F*_(3.851, 130.9)_ = 78.78, *p* < 0.0001), as well as a main effect of housing condition (*F*_(1, 34)_ = 4.17, *p* = 0.048) but no main effect of sex (*F*_(1, 34)_ = 0.069, *p* = 0.793). There was no interaction between time and sex (*F*_(5, 170)_ = 1.15, *p* = 0.336), time and housing condition (*F*_(5, 170)_ = 1.26, *p* = 0.285), or sex and housing condition (*F*_(1, 34)_ = 0.153, *p* = 0.697), nor was there a significant three-way interaction between these variables (*F*_(5, 170)_ = 0.669, *p* = 0.646). Post-hoc analysis did not reveal any significant differences in freezing behavior across groups at any time point (adjusted *p* > 0.05).

### Home Cage Ethanol Drinking

As previous studies in rodents have demonstrated that adolescent social isolation increases home cage ethanol self-administration (McCool and Chappell, 2009, Butler et al., 2014a, Skelly et al., 2015, Butler et al., 2016), we next assessed whether adolescent social isolation affects binge ethanol drinking in male and female C57BL/6J mice across time using a modified version of the standard DID paradigm that allowed us to assess ethanol preference on day 4 of each DID cycle (**Figure 3A,D**; n = 10 per group). A mixed-effects analysis was used to evaluate consumption of 20% ethanol across four cycles in GH and SI females **(Figure 3A**, left), revealing a main effect of cycle (*F*_(6.372, 112.6)_ = 4.32, *p* < 0.0001) but no main effect of housing condition (*F*_(1, 18)_ = 1.24, *p* = 0.280) or interaction between these variables (*F*_(15, 265)_ = 1.43, *p* = 0.132). To ensure that a group difference was not being obscured by a ceiling effect, we next increased the concentration of ethanol to 30% for two cycles, and a mixed-effects analysis revealed no effects or interactions at this concentration either (main effect of cycle: *F*_(3.621, 60.52)_ = 1.77, *p* = 0.153; main effect of housing condition: *F*_(1, 17)_ = 0.219, *p* = 0.645; time by housing condition interaction: *F*_(7, 117)_ = 1.72, *p* = 0.111). We also found no effect of social isolation on ethanol preference at either concentration in females (**Figure 3A**, right). A mixed-effects analysis of 20% ethanol preference revealed no effects (main effect of cycle: *F*_(3, 49)_ = 0.097, *p* = 0.961; main effect of housing condition: *F*_(1, 18)_ = 1.71, *p* = 0.207; cycle by housing condition interaction: *F*_(3, 49)_ = 2.19, *p* = 0.101). Similarly, a two-way RM ANOVA assessing 30% ethanol preference revealed no effects (main effect of time: *F*_(1, 17)_ = 1.07, *p* = 0.316; main effect of housing condition: *F*_(1, 17)_ = 3.83, *p* = 0.252; time by housing condition interaction: *F*_(1, 17)_ = 1.83, *p* = 0.194).

**Figure 3.**
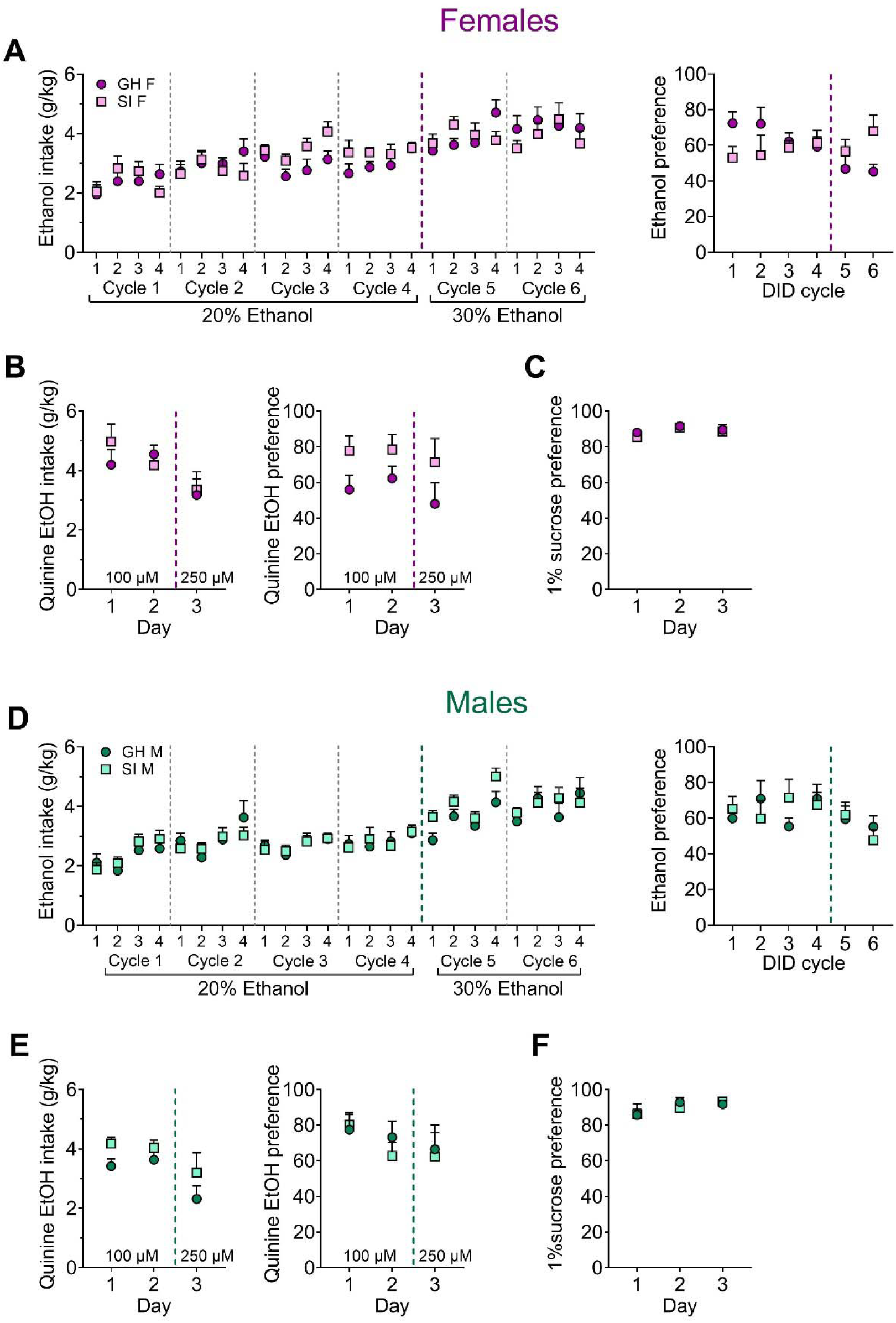
Effects of adolescent social isolation on home cage ethanol drinking and reward and aversion sensitivity in adult female (A-C) and male (D-F) mice. (**A**) There are no effects of adolescent SI on binge ethanol consumption (left) or 24-hr ethanol preference (right) across six weeks of 20% and 30% ethanol in a modified EtOH DID paradigm in females. (**B**) Adolescent GH and SI females display similar consumption of quinine-adulterated ethanol (left) and preference for it over water (right) across multiple quinine concentrations. (**C**) Adolescent SI does not alter preference for a 1% sucrose solution over water in female mice. (**D-F**) Similarly, adolescent SI in males does not alter ethanol intake or preference (**D**), quinine-adulterated ethanol intake or preference (**E**), or 1% sucrose preference (**F**).

Similar to females, social isolation did not affect ethanol consumption or preference in males (**Figure 3D**, left). A mixed-effects analysis of 20% ethanol consumption (**Figure 3D**; n = 10 per group) revealed a significant main effect of cycle (*F*_(7.450, 132.6)_ = 4.10, *p* < 0.001), but no main effect of housing condition (*F*_(1, 18)_ = 0.004, *p* = 0.947) or interaction between these factors (*F*_(15,_ 267) = 0.527, *p* = 0.924). A mixed-effects analysis of 30% ethanol intake also revealed a main effect of cycle (*F*_(7, 121)_ = 7.36, *p* < 0.001), but no main effect of housing condition (*F*_(1, 18)_ = 1.29, *p* = 0.270) or significant cycle by housing condition interaction (*F*_(7, 121)_ = 1.63, *p* = 0.132). A mixed-effects analysis of 20% ethanol preference (**Figure 3D**, right) compared to water revealed no effects (main effect of cycle: *F*_(2.357, 38.49)_ = 0.325, *p* = 0.758; main effect of housing condition: *F*_(1, 18)_ = 0.213, *p* = 0.649; cycle by housing condition interaction: (*F*_(3, 49)_ = 2.06, *p* = 0.117). Similarly, a mixed effects analysis assessing 30% ethanol preference did not reveal significant group differences (main effect of cycle: *F*_(1, 35)_ = 1.88, *p* = 0.179; main effect of housing condition: *F*_(1, 35)_ = 0.151, *p* = 0.699; cycle by housing condition interaction: *F*_(1, 35)_ = 0.536, *p* = 0.468).

### Aversion-Resistant Ethanol Drinking

To assess whether adolescent social isolation alter aversion-resistant ethanol consumption, we measured home cage DID intake using 20% ethanol adulterated with quinine (**Figure 3B,E**). Mice were given 4 hr access to 20% ethanol containing either 100µM quinine (days 1 and 2, average used for analysis) or 250µM quinine (day 3). Among female mice (ns = 9), a two-way RM ANOVA for quinine-adulterated ethanol intake did not reveal any significant differences (**Figure 3B**; main effect of quinine concentration: *F*_(1, 16)_ = 4.21, *p* = 0.056; main effect of housing condition: *F*_(1, 16)_ = 0.175, *p* = 0.681; concentration by housing condition interaction: *F*_(1,_ 16) = 0.001, *p* = 0.977). Similarly, a two-way RM ANOVA assessing quinine-adulterated ethanol preference revealed no main effects of housing condition (*F*_(1, 16)_ = 3.62, *p* = 0.074) or quinine concentration (*F*_(1, 16)_ = 1.67, *p* = 0.214), nor any significant interaction between these variables (*F*_(1, 16)_ = 0.049, *p* = 0.826). In male mice (GH n = 10, SI n = 9), there was a significant main effect of quinine concentration on ethanol intake (**Figure 3E**; *F*_(1, 17)_ = 2.93, *p* = 0.105), with the higher dose of quinine suppressing ethanol consumption. However, there was no significant main effect of housing condition (*F*_(1, 17)_ = 2.93, *p* = 0.105), nor a significant interaction between these factors (*F*_(1, 17)_ = 0.128, *p* = 0.724). A two-way RM ANOVA comparing ethanol preference across quinine concentrations did not reveal any significant differences between GH and SI male mice (main effect of quinine concentration: *F*_(1, 17)_ = 1.29, *p* = 0.271; main effect of housing condition: *F*_(1, 17)_ = 0.108, *p* = 0.746; concentration by housing condition interaction: *F*_(1, 17)_ = 0.001, *p* = 0.981).

### Sucrose Preference Test

To determine whether social isolation during adolescence impacts general reward sensitivity, we measured 1% (w/v) sucrose preference versus water across three days (**Figure 3C,F**). A two-way RM ANOVA comparing adolescent GH (n = 10) and SI (n = 9) female mice revealed a significant main effect of time (**Figure 3C**; *F*_(1.687, 28.68)_ = 4.32, *p* = 0.028) but no main effect of housing condition (*F*_(1, 17)_ = 0.342, *p* = 0.566) or interaction between these variables (*F*_(2, 34)_ = 0.255, *p* = 0.775). In male mice, no differences in sucrose preference emerged (**Figure 3F**; main effect of time: *F*_(1.418, 25.53)_ = 2.57, *p* = 0.110; main effect of housing condition: *F*_(1, 18)_ = 0.025, *p* = 0.874; time by housing condition interaction: *F*_(2, 36)_ = 0.331, *p* = 0.720). Altogether, results from our drinking experiments suggest that binge ethanol consumption, aversion-resistant ethanol intake, and general reward sensitivity were unaltered by adolescent social isolation.

### Home Cage Social Interaction

We found a robust effect of adolescent, but not adult, social isolation on increased social behavior in adulthood using a social interaction paradigm in a novel environment. We further probed the stability and generalizability of this phenotype using a home cage social interaction test in which the experimental mouse remained in its home cage and a novel intruder conspecific was placed in the cage for five min (**Figure 4**; GH females n = 9, SI females n = 9, GH males n = 10, SI males n = 9). Adolescent SI males and females again showed greater social interaction in this paradigm. A two-way ANOVA on the total number of social interaction bouts (**Figure 4A**) showed a main effect of housing condition (*F*_(1, 33)_ = 19.08, *p* = 0.0001) and no effect of sex (*F*_(1,_ 33) = 3.99, *p* = 0.054) or sex by housing interaction (*F*_(1, 33)_ = 1.06, *p* = 0.310). Post-hoc t-tests confirmed this effect occurred in both females (*t*_(33)_ = 2.33, adjusted *p* = 0.026) and males (*t*_(33)_ = 3.87, adjusted *p* = 0.001). This increased interaction was true for both head-to-head and head-to-tail interactions. Head-to-head (**Figure 4B**): main effect of housing (*F*_(1, 33)_ = 9.78, *p* = 0.004), no effect of sex (*F*_(1, 33)_ = 4.13, *p* = 0.050), and no interaction (*F*_(1, 33)_ = 0.002, *p* = 0.961); post-hoc t-tests: *ps* > 0,05. Head-to-tail (**Figure 4C**): main effect of housing (*F*_(1, 33)_ = 16.26, *p* = 0.0003), no effect of sex (*F*_(1, 33)_ = 1.72, *p* = 0.198), and no interaction (*F*_(1, 33)_ = 2.43, *p* = 0.128); post-hoc t-tests showed the effect was driven by males: females (*t*_(33)_ = 1.73, adjusted *p* = 0.094), males (*t*_(33)_ = 4.01, adjusted *p* = 0.0007). In contrast to social interactions, there was no effect of adolescent SI on digging or climbing behaviors (**Figure 4D,E**). Two-way ANOVAs on the number of digging bouts and the number of climbing bouts showed no effects of housing condition, sex, or an interaction (*ps* > 0.05).

**Figure 4.**
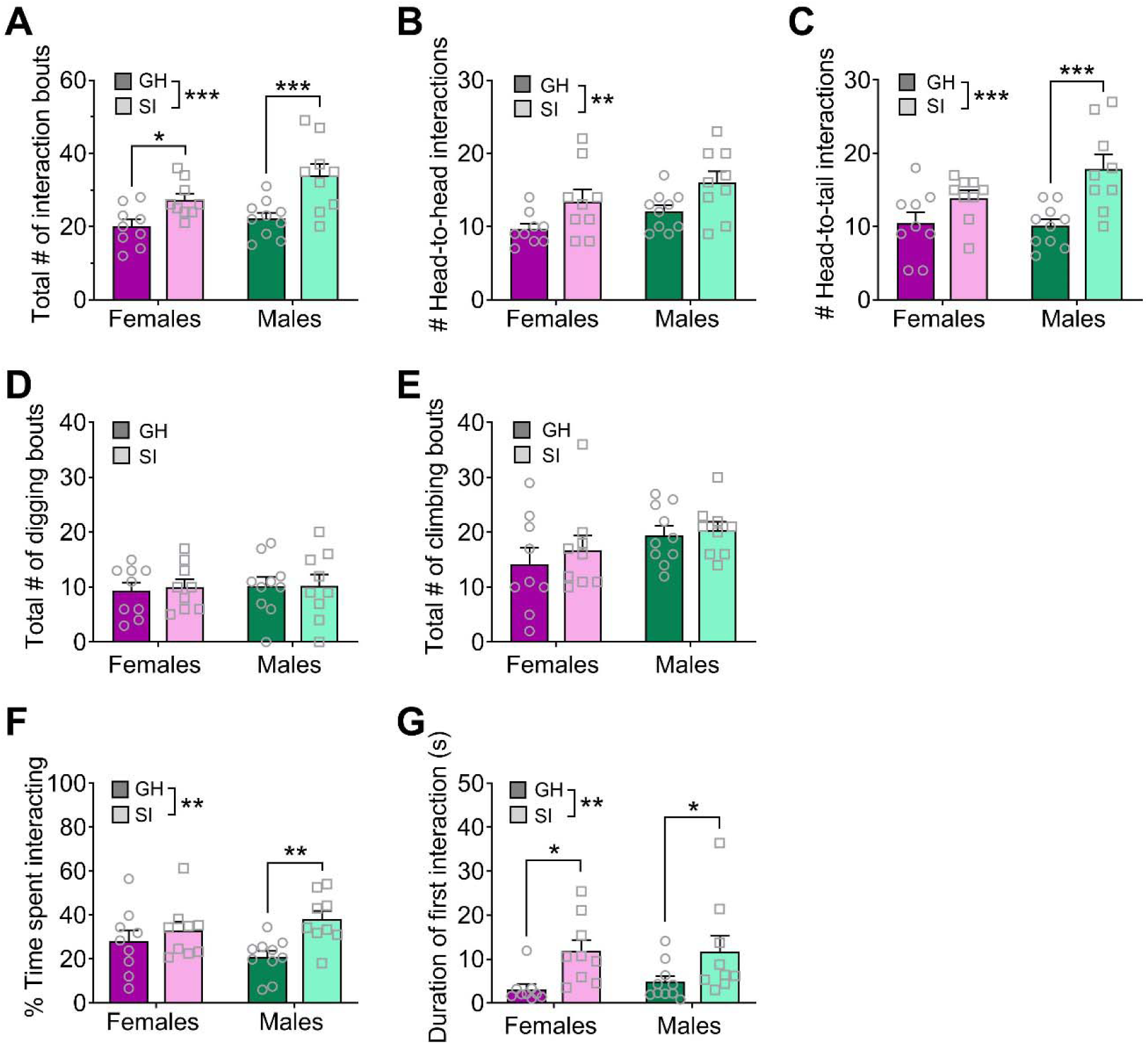
Effects of adolescent social isolation on home cage social interaction in adulthood. (**A**) aSI mice display an increased number of social interaction bouts in both males and females. (**B-C**) This overall phenotype is present when only head-to-head interactions (**B**) or head-to-tail interactions (**C**) are considered. (**D-E**) In contrast, digging (**D**) and climbing (**E**) behaviors are not altered by adolescent SI. (**F**) Adolescent SI mice spend a greater proportion of the 5 min assay interacting with the stranger mouse than their GH counterparts, and effect driven by males. (**G**) The duration of the first social interaction bout is longer in adolescent SI mice of both sexes. Data are expressed as means + SEM; \**p* < 0.05, \*\**p* < 0.01, \*\*\**p* < 0.001 between groups.

Given this distribution of behaviors during the home cage assay, adolescent SI mice spent a greater proportion of time engaged in social interaction than their GH counterparts (**Figure 4F**). A two-way ANOVA on the percent time spent exploring a novel social partner revealed a significant main effect of housing (*F*_(1, 33)_ = 7.59, *p* = 0.010) but no main effect of sex (*F*_(1, 33)_ = 0.055, *p* = 0.815) or interaction between these variables (*F*_(1, 33)_ = 2.44, *p* = 0.127). Post-hoc analysis showed that the effect of social isolation was driven by males (*t*_(33)_ = 3.09, adjusted *p* = 0.008) but did not occur in females. However, the duration of the first interaction bout was longer in adolescent SI mice of both sexes (**Figure 4G**). A two-way ANOVA assessing the duration of the first bout of social interaction revealed a significant main effect of housing condition (*F*_(1, 33)_ = 11.23, *p* = 0.002), but no main effect of sex (*F*_(1, 33)_ = 0.109, *p* = 0.742) or significant sex by housing condition interaction (*F*_(1, 33)_ = 0.143, *p* = 0.707). Post-hoc analysis confirmed that both SI females and males spent more time interacting with a novel social partner during this first bout than their GH counterparts (females: *t*_(33)_ = 2.61, adjusted *p* = 0.027; males: *t*_(33)_ = 2.13, adjusted *p* = 0.040). Interestingly, however, SI mice had a longer latency to first approach the stranger mouse, suggesting some initial inhibition of this hypersocial behavior (data not shown). A two-way ANOVA revealed a main effect of housing condition (*F*_(1, 33)_ = 19.00, *p* < 0.001), but no main effect of sex (*F*_(1, 33)_ = 2.06, *p* = 0.160) or sex by housing interaction (*F*_(1,_ 33) = 1.35, *p* = 0.254). Post-hoc analysis revealed that SI males and females took significantly more time to approach the novel social partner than their GH counterparts (GH females vs SI females: *t*_(33)_ = 2.23, adjusted *p* = 0.039; GH males vs SI males: *t*_(33)_ = 3.95, adjusted *p* = 0.001). In spite of this initial delay in interaction, the overall results support our initial findings that adolescent social isolation produces an aberrant hyper-social phenotype in adulthood in C57BL/6J mice.

## Discussion

These studies were designed to assess whether the harmful and translationally-relevant behavioral consequences of adolescent SI well-characterized in rats can be reliably recapitulated in C57BL/6J mice, the most common laboratory mouse background strain. We further sought to determine whether adolescence is a critical period for behavioral plasticity or whether a similar long-term social isolation in adulthood impacts these pathology-related behaviors. Surprisingly, we did not see any consistent phenotypes following adult SI, as mice displayed an anxiogenic phenotype in the light/dark box assay (**Figure 2E**) but not on any other measures of anxiety-like behavior. These findings indicate that singly housing mice in adulthood, as is done routinely in alcohol and drug self-administration studies, among others, does not alter basal behavioral states in C57BL/6J mice; thus, adult isolation is not a major confounding variable for most behavioral assays including those measured herein. Similarly, we found few effects of adolescent social isolation on performance in a battery of behaviors, which was surprising given the literature showing the deleterious effects of stress during the adolescent period on adult behaviors. However, the most robust effect of adolescent social isolation we observed was that it promoted social behavior in adulthood in both sexes (**Figure 1D**; **Figure 4**), and effect remarkably similar in nature to the stress imposed upon the mice.

Contrary to our predictions, we did not find that adolescent social isolation increases anxiety-like behavior in male or female C57BL/6J mice (**Figure 1**). In fact, following adolescent isolation, adult female mice spent more time in the open arms of the elevated plus maze on average, a behavior which is classically interpreted as a sign of anxiolysis (**Figure 1B**). This anxiolytic effect of adolescent isolation in mice has been reported elsewhere (Voikar et al., 2005, Lopez and Laber, 2015). Previous studies have also found some evidence that adolescent social isolation induces an anxiogenic phenotype in the light/dark box and hyperlocomotion in the open field test in mice (Voikar et al., 2005, Gan et al., 2014, Amiri et al., 2015, Medendorp et al., 2018), but these results have not always been reported (Koike et al., 2009). In contrast to, we found no effect of adult social isolation on anxiety-like behavior in the EPM (**Figure 2B**), suggesting some adolescent time period specificity for this effect. Intriguingly, we found that adult social isolation increased anxiety-like behavior in the light/dark box, suggesting that if anything, adult isolation produces the opposite effect of adolescent isolation. However, in both cohorts, other measures of anxiety-like behavior did not recapitulate these effects, suggesting there are no reliable effects of social isolation at either time point on adult anxiety-related behavior in C57BL/6J mice.

Perhaps our most striking finding is that isolation rearing during adolescence increased social exploration and interaction in adulthood. Specifically, we found that preference for a novel social partner increased in both males and females following protracted adolescent isolation (**Figure 1D**). We extended this finding in a home cage social interaction test with a novel intruding conspecific (**Figure 4**), demonstrating that this hypersocial behavior occurs in both familiar and novel environments. Aberrantly high social exploration may be maladaptive in settings in which social caution or defensive behavior is more appropriate, such as during exposure to an unfamiliar intruder. This phenotype is similar to that observed in some developmental disorders such as Williams’ Syndrome, in which individuals inappropriately approach and engage with strangers. However, as this behavior occurred following a longer delay before approaching the stranger mouse, the social phenotype of the adolescent SI mice could be a compensatory mechanism that actually promotes an adaptive social phenotype beneficial in certain contexts that require social affiliation for survival. This pro-social interpretation has previously been reported to occur in female mice following exposure to a developmental stressor (Koike et al., 2009, Bondar et al., 2018). Interestingly, many groups have reported the exact opposite effect of adolescent isolation on social behavior in mice, finding that this developmental stressor decreases social interest in adulthood (Balemans et al., 2010, Medendorp et al., 2018). Nonetheless, reduced social learning (Kercmar et al., 2011) and aberrant social behavior when placed back into group housing in adulthood (Endo et al., 2018) have also been reported following post-weaning isolation in C57BL/6J male and female mice, further supporting a specific role for peri-adolescent social isolation in abnormal adult social behavior. This is unsurprising given that this is a crucial developmental period for the development of prosocial behaviors (Spear, 2004, Panksepp et al., 2007, Panksepp and Lahvis, 2007).

Interestingly, we did not identify a robust effect of isolation in adulthood on measures of social interaction (**Figure 2D**), further suggesting that adolescence is a critical period for the development of sensitivity to social reward. We also tested interest in a non-social novel object following adolescent social isolation and found no significant effect of rearing condition on novel object preference (**Figure 1F**). Again, no differences in novel object preference emerged following social isolation in adulthood, although GH males spent more total time exploring the social partner and novel object combined than SI males or GH females (**Figure 2F**).

Prolonged social isolation during adolescence or adulthood has also been reported to impact aspects of fear memory formation in rats and mice (Pibiri et al., 2008, Pinna et al., 2008, Lukkes et al., 2009a, Okada et al., 2015, Pinna, 2019). Here we tested the effect of sex and housing condition on fear learning across six tone/footshock pairings. We did not identify any effect of housing condition on fear memory formation following adolescent isolation (**Figure 1G**) but did observe delayed acquisition following adult isolation, however final acquisition was similar across all groups (**Figure 2G**). Together, these results suggest that singly housing C57BL/6J mice during adolescence or adulthood does not reliably impact fear memory formation.

Adolescent isolation has been demonstrated to increase alcohol self-administration in male rats and both male and female mice (Lopez et al., 2011, Butler et al., 2014b, Lopez and Laber, 2015, Skelly et al., 2015, Butler et al., 2016). Here, we evaluated adolescent social isolation on binge alcohol drinking using a modified DID paradigm and found no effects on 20%, 30%, and quinine-adulterated 20% ethanol consumption or preference, nor on a rewarding 1% sucrose solution, in either sex (**Figure 3**). Our results are inconsistent with the findings of Lopez and colleagues (Lopez and Laber, 2015), who found that adolescent social isolation in C57 mice produced a small but significant increase in alcohol consumption at one time point. However, that study did not exam chronic home cage ethanol self-administration. Regardless, our data indicate that perhaps the effects of chronic social stress in adolescence on ethanol drinking are less robust than the effects reported in rats. Interestingly, adolescent social isolation has been reported to produce a protracted increase in ethanol intake and preference in male C57BL/6J mice given intermittent access to ethanol in their home cage, but only at a relatively low ethanol concentration (5%); these differences disappeared when animals were offered a higher concentration of ethanol (20%) (Advani et al., 2007). Together, these findings generally suggest that adolescent isolation does not reliably produce a translationally relevant escalation of ethanol self-administration in C57BL/6J mice.

In general, we found that C57BL/6J mice are not reliably sensitive to isolation stress. Beyond the findings outlined herein, others have presented some evidence that single housing may not be experienced as an adversity among C57BL/6J mice (Bartolomucci et al., 2003, Arndt et al., 2009), and in fact may actually decrease social stress in males of this species (Singewald et al., 2009). Others have not found evidence to support a protective effect of adolescent social isolation in female C57BL/6J mice (Martin and Brown, 2010). Interestingly, the majority of studies reporting a behavioral effect of adolescent isolation on anxiety-like behavior, fear memory formation, or drug self-administration in have initiated isolation at the same time that play behavior is typically increasing, suggesting that disruption of play behavior may be a major contributor to this phenotype (Walker et al., 2019). As mice engage in less social play in adolescence than rats, this may partly explain the variability in the behavioral effects of adolescent isolation rearing reported here and elsewhere. Although these findings present an issue for researchers interested in identifying the link between developmental stress and psychopathology using mouse models on a C57BL/6J strain, the most common background for genetic manipulation, it also suggests that experimentally-mandated individual housing in adolescence or adulthood may not produce confounding effects on basal behavioral states that experimenters prefer to avoid.

## Author contributions

JKRI and MJS collected and analyzed the data. JKRI, MJS, and KEP designed the studies and wrote and edited the manuscript.

